# Heavy metal pollution impacts soil bacterial community structure and antimicrobial resistance at the Birmingham 35^th^ Avenue Superfund Site

**DOI:** 10.1101/2022.04.12.488090

**Authors:** Anuradha Goswami, Sarah J. Adkins-Jablonsky, Marcelo Malisano Barreto Filho, Michelle D. Schilling, Alex Dawson, Sabrina Heiser, Aisha O’Connor, Melissa Walker, Qutia Roberts, J. Jeffrey Morris

## Abstract

Heavy metals (HMs) are known to modify bacterial communities both in the laboratory and *in situ*. As a consequence, soils in HM contaminated sites like the U.S. Environmental Protection Agency (EPA) Superfund sites are predicted to have altered ecosystem functioning, with potential ramifications for the health of organisms, including humans, that live nearby. Further, several studies have shown that metal tolerant bacteria are often also resistant to antimicrobial agents (AMR), and therefore HM contaminated soils could potentially act as reservoirs that could disseminate AMR genes into human-associated pathogenic bacteria. To explore this possibility soil samples were collected from six public locations in the zip code 35207 (the home of the North Birmingham 35^th^ Avenue Superfund site) and in six public areas in a neighboring reference zip code (35214). Sequencing of the V4 region of the bacterial 16S rRNA gene revealed that elevated concentrations of HMs As, Mn, Pb, and Zn reduced microbial diversity and altered community structure within each zip code. While there was no difference between zip codes in the proportion of total culturable microbes that survived antimicrobial or metal exposure, bacterial isolates with HMR almost always also exhibited AMR. Metagenomes inferred using PICRUST2 also predicted significantly higher mean relative frequencies in 35207 for several AMR genes related to both specific and broad-spectrum AMR phenotypes. Together, these results support the hypothesis that chronic HM pollution alters soil bacterial community structure in ecologically meaningful ways and may also select for bacteria with increased potential to contribute to AMR in human bacterial disease.

## INTRODUCTION

Heavy metals (HMs) are necessary for biological processes across all domains of life (e.g. by acting as catalytic cofactors in proteins) but they can also be toxic in high concentrations. HMs in soil environments are documented human health risks (Zhao et al. 2012), a fact all too familiar to people in the community of North Birmingham. For decades, environmental injustice in the form of industrial pollution from coke furnaces and steel plants has plagued residents living in this Central Alabama area, over 90% of whom are African-American and 40% of whom live under the federal poverty line (Allen et al., 2019). These facilities emit particulate matter containing HMs like Cd, As, and Mn into the air and soil. In recognition of the potential health impacts caused by this large-scale pollution, the Environmental Protection Agency (EPA) designated North Birmingham as the 35th Avenue Superfund Site in 2012 (henceforth referred to by its zip code, 35207), committing the U.S. federal government to fund pollution cleanup (Allen et al., 2019). However, continued pollution and local politics have stalled EPA’s progress, and 35207 residents have yet to see substantial progress towards confronting and overcoming the legacy of environmental mismanagement. Grassroots organizations such as People Against Neighborhood Industrial Contamination (PANIC) have asked for remediation and financial restitution for 35207 residents, but also more scientific studies to understand and quantify the environmental and health impacts explicitly related to HM contamination in their community (GASP, 2014). In the words of Charlie Powell, the founder of PANIC: “We been fighting this for years and we ain’t been getting no justice.” (Hodgin, 2020)

Chronic HM exposure is known to have several effects on microbial communities. HM pollution can select for specific bacterial taxa and physiological properties (Kandeler et al., 1996), which can in turn impact the diversity of the microbial populations (Epelde et al., 2015; Kandeler et al., 1996). For instance, HMs can affect soil properties, like spatial structure, in ways that in turn increase bacterial community diversity (Bourceret et al. 2016; Rajeev et al., 2020; Thomas et al., 2020). HMs are also known to cross-select for both heavy metal resistance (HMR) and broad antibiotic/antimicrobial resistance (AMR) in bacteria (Baker-Austin et al., 2006; Seiler and Berendonk, 2012), even at low levels (Gullberg et al., 2014), because many of the mechanisms conferring resistance to one set of toxins are also effective at resisting the other as well (Baker-Austin, Wright et al. 2006). Laboratory experiments *in vivo* showed, for example, that cadmium exposure induced transmembrane efflux pump resistance to not only HMs like Zn and Cd, but also to carbapenem antibiotics (Perron et al., 2004).

Thus, it is possible that 35207 soils contain microbial communities not only with different microbial populations than surrounding soils, but also containing bacteria with high levels of AMR. The development of bacterial cross-resistance between heavy metal and antibiotic genes (Chen et al., 2019; Timoney et al. 1978) can occur through horizontal gene transfer (Schlüter et al., 2008; Szczepanowski et al., 2008; Li et al., 2015), and therefore it is possible that these bacteria could function as a reservoir from which AMR could spread to the human-associated bacteria of 35207 residents, creating an additional health risk beyond the direct impacts of HM exposure. Alarmingly, many AMR genes confer broad resistance across antibiotic classes (Blair et al., 2015). According to the World Health Organization, the spread of AMR is one of the most pressing concerns for the 21^st^ century (World Health Organization, 2014). Given that 35207 residents are already more vulnerable to infectious diseases (Dyer, 2020) and that AMR infections continue to rise globally (Levy, 2004), investigating the potential for industrial HM pollution to favor AMR at sites like 35207 is a social justice imperative. In order to investigate this possibility, we collected soil samples from 35207 as well as from a nearby Birmingham neighborhood with similar demographics but farther from the EPA-designated Superfund site (henceforth also referred to by its zip code, 35214) and used them to address the following questions:

1. How do soil bacterial communities from the HM-polluted neighborhood differ from those in a lesser polluted neighborhood?
2. Do HM-polluted samples show physiological or predictive genetic evidence of AMR?

## METHODS

### Site description and soil sampling

This project began as a set of course-based undergraduate research experiences using methods as previously described (Adkins-Jablonsky et al., 2020). In August 2019, three soil samples each were collected at six different public access parks in the North Birmingham 35207 zip code, for a total of eighteen soil samples. Each sample (5g) was collected from the topsoil layer with an ethanol-sanitized trowel and a sterile plastic bag. Sampling was repeated in the 35214 zip code for 18 additional soil samples. All three samples per park were then homogenized by vigorous shaking, for a total of 12 homogenized soil samples (∼15g each), six from 35207, and six from 35214 (Fig. 1A). To calculate percent organic mass via percent weight loss-on-ignition (LOI), samples were oven dried at 100°C for 1 hour, weighed, heated in a muffle furnace at 400°C for 3 hours, and re-weighed. pH was measured via electrode after suspending 1g of soil in 2.5 mL distilled water. Samples were stored at −80°C until further processing. In September 2020, soil sampling was repeated from five of the original six parks per zip code to provide fresh soil cultures for culture assays. One public park per zip code was not re-sampled due to access limitations during the COVID-19 pandemic.

**Figure 1.**
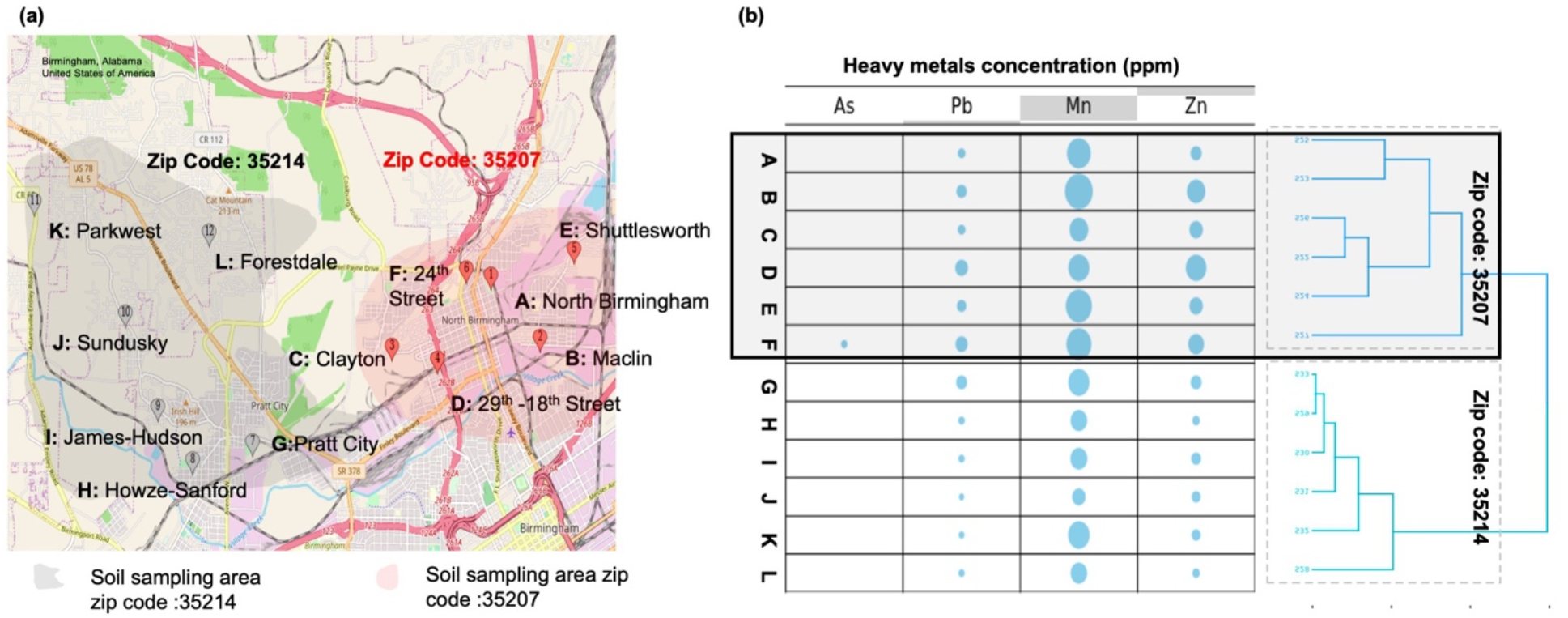
**(a)** Map of sampling sites from G-to-K numbered 7-12 from 35214 (in grey) and A-to-F from 32507 (in red). Map generated by GoogleMaps ©. Map Customizer with longitude and latitude points is accessible at https://www.mapcustomizer.com/map/Nbham%20paper; **(b)** Relative metal concentrations are depicted in the bubble chart. Metal analytes expressed “0” indicates Below Detection Limit (BDL) based on EPA Method 6010B (EPA laboratory ID AL01084). Cd not shown as Cd analytes were BDL. The dendrogram on right of the chart shows the k-mers clustering of sampling sites, clustering sites based on heavy metals (HMs) levels.

### Heavy metal testing

Soil (56 g from each 2019 sample) was tested for heavy metal concentrations by Sutherland Environmental Testing using EPA Method 6010B (EPA laboratory ID AL01084) through Inductively Coupled Plasma-Atomic Emission Spectroscopy. Analytes reported were As, Cd, Pb, Mn, and Zn. The estimated instrumental detection limits of As, Cd, Pb, Mn and Zn according to EPA Method 6010B were 35, 2.3, 28, 0.93 and 1.2 μg/L, respectively. Estimated concentrations below these detection limits were assumed to be 0 for the purposes of our analyses. K-mer clustering of collection sites based on their HM profiles was done using the *kmer* package (Wilkinson, 2018) in R v1.4.1717 (RStudio Team, 2020).

### Nucleic Acid Purification and Sequencing

Bacterial genomic DNA was extracted from 0.20 g of soil from each of the 2019 collections using the Qiagen Dneasy PowerSoil Pro kit (Germantown, MD, USA, Cat No./ID: 47016) according to the manufacturer’s instructions. Presence and quality of DNA was confirmed by spectrophotometry using Gen5 software with a Take3 Micro-Volume Plate in a BioTek Synergy H1 microplate reader (A_260/280_ ∼1.8; each sample contained at least 65 ng/μL of DNA). At the University of Alabama at Birmingham Heflin Center for Genomics (Birmingham, Alabama), an amplicon library was created via PCR amplification of the hypervariable region 4 (V4) of the 16S rRNA gene using barcoded oligonucleotide primers F515 (CACGGTCGKCGGCGCCATT) and R806 (GGACTACHVGGGTWTCTAAT) (Caporaso et al., 2011). Genomic DNA was then gel-purified and sequenced using the Illumina MiSeq platform.

### Microbiome sequence processing and statistical analysis

DNA sequences were processed with the Quantitative Insights Into Microbial Ecology package (qiime2-2020.11) (Bolyen et al., 2019; Caporaso et al., 2010) (Table S1). The demultiplexed paired end reads obtained from Illumina sequencing were processed with the denoising algorithm DADA2 (Callahan et al., 2016). The minimum and maximum demultiplexed read counts were 163408 and 359799 respectively. The sequences were truncated to 250 base pairs length. The representative sequences obtained after denoising were clustered using q2-vsearch (QIIME2 plugin) denovo clustering at 97% similarity threshold to remove singletons and obtain 4185 unique operational taxonomic units (OTUs). The OTUs were classified using a pre-trained Naive Bayes classifier on silva-138-99-515-806-nb-classifier, earlier trained on 515F/806R region of sequences from the Silva-138 99% OTUs database (QIIME2 Data resources - MD5: e05afad0fe87542704be96ff483824d4 (Quast et al., 2013; Bokulich et al., 2018), link: https://docs.qiime2.org/2021.2/data-resources) (Table S2). The q2-feature-classifier plugin was used to taxonomically identify OTUs and remove non-bacterial OTUs (Bokulich et al., 2018).

Alpha diversity (Shannon and Simpson indices) at genus level was estimated using *MicrobiomeAnalyst* (https://www.microbiomeanalyst.ca) (Chong et al., 2020, Dhariwal et al., 2017) after applying filters for low count (<4 counts in more than 20% of samples) and low variance (limited to the interquartile range) features and normalizing by total sum scaling. For beta-diversity analyses, the OTU table was first rarefied to the level of the least deeply sequenced sample using a ranked subsampling algorithm (Beule and Karlovsky 2020). Canonical correspondence analysis (CCA) was completed using the VEGAN v2.5 package (Dixon, 2003) in R v1.4.1717 (RStudio Team, 2020). Principle coordinates and NMDS ordination were performed in mothur (Schloss, Westcott et al. 2009) using distance matrices generated with one of several different algorithms; the ordination technique and distance method that gave the best r-squared value on 3 axes (NMDS with the Yue-Clayton theta metric, r-squared = 0.97, stress = 0.065) was retained for further analysis. Statistical significance of the impact of environmental metadata on NMDS ordination was assessed by calculating the Spearman correlation coefficient between each variable and the first two NMDS axes using the corr.axes command in mothur (Table S3). The influence of individual OTUs on NMDS ordination was also determined with corr.axes (Table S2).

Differential abundance of OTUs between zip codes was determined using Metastats (implemented through mothur) (Paulson, Pop et al. 2011) and LEFSe (implemented through MicrobiomeAnalyst) (Segata, Izard et al. 2011) (Table S2). Spearman correlations between OTU (Table S2) and higher taxon abundances (Table S4) and environmental metadata (metals, pH, organic carbon) were calculated in R. Significance levels of differences in OTU abundances between zip codes was computed using a simple 2-tail t-test for each OTU individually (Table S4). Only OTUs that represented at least 1% of the overall community in at least one sample were considered further for these OTU-level analyses.

### Predictive metagenomic analysis

16S amplicon data was used to predict microbial population metagenomes using Phylogenetic Investigation of Communities by Reconstruction of Unobserved States (PICRUSt v2.3.0) (Langille et al., 2013) (Table S6). The 16S rRNA sequence and abundance *biom (Biological Observation Matrix)* table was used to predict Kyoto Encyclopedia of Genes and Genomes (KEGG) genes and pathway abundances. Sequences with Nearest Sequence Taxon ID (NSTI) above a cut-off of 2.0 were removed along with their sequence counts (Table S5). The OTU abundances were multiplied by the corresponding NSTI value of each OTU and weighted NSTIs of each sample were calculated from the sum of the column per sample and divided by the total read depth per sample (0.27 ± 0.02 from 35207 and 0.25 ± 0.035 from 35214). Weighted NSTI values describe the degree to which microbes in samples are related to known genomes, where a value of 0.10 would represent 90% representational similarity. While lower numbers are preferred, NSTI values of up to 0.30 have been recorded for microbial populations derived from soil samples and yielded useful representations of shotgun metagenomes (Langille, Zaneveld et al. 2013). The predicted metagenomes were functionally annotated with PICRUSt2 using the KEGG pathway database (Kanehisa and Goto, 2000). The KO IDs obtained were manually annotated using the KEGG database to estimate AMR and HMR gene abundance. KEGG genes were compared between the two zip codes using Welch’s T test in STAMP v2.1.3 (Statistical analysis of taxonomic and functional profiles) Bioinformatics software (Parks et al 2014). The relationship between gene abundances and environmental metadata was computed as a linear model in R, as was the Fisher’s Exact Test to determine if AMR or HMR genes were more likely to be significantly correlated with metals than other genes. Over-representation of KEGG pathways between the zip codes was estimated using Over Representation Analysis (ORA) with the command *enrichKEGG* in clusterProfiler (Table S7).

### Microbial cultivation, identification, and antimicrobial sensitivity determination

Microbial cultures were isolated on PYT80 agar containing (per L) 80 mg each of peptone, yeast extract, and tryptone, 1.95 g 2-(N-morpholino)ethanesulfonic acid, 15 g of purified agar (USP Grade, MP Biomedicals), and 10 mg cycloheximide and adjusted to pH 6.50 (modified by Lin et al., 2012). 1 mg of soil from each 2020 sample was suspended in 9 mL 0.085% sterile saline and then serially diluted onto PYT80 agar with or without additional heavy metals (0.5M Pb(NO_3_)_2_ at a final concentration of 0.4mM; 1M MnCl_2_ at a final concentration of 25mM; or 0.5M ZnCl_2_ at a final concentration of 0.5mM) or antibiotics (erythromycin at a final concentration of 50 μg/mL; ampicillin at a final concentration of 100 μg/mL; or kanamycin at a final concentration of 25 μg/mL). After inoculation, plates were incubated at 20°C for a minimum of 72 hr. The overall community sensitivity to the amendments was calculated as the log fold change of the ratio of CFU/mL of HMR or AMR resistant culturable organisms to total culturable organisms (determined by plate counts on amended vs. unamended PYT80 plates, respectively).

We also isolated a total of 46 HMR bacterial strains that could grow on PYT80 supplemented with Mn, Zn, or Pb, including representatives from all 12 sites, and identified them by PCR amplification of the 16S rRNA gene using primers UA1406R (5’-ACGGGCGGTGWGTRCAA-3’) and U341F (5’-CCTACGGGRSGCAGCAG-3’) followed by Sanger sequencing at the UAB Heflin Center for Genomics (Birmingham, Alabama). Sequences were trimmed using MEGA X (v.10.2.4) (Kumar et al., 2018), and identified using the Basic Local Alignment Search Tool (BLAST) 16S/ITS BLASTn function against the nr database (BLAST 2.11.0) (Altschul et al., 1990) (Table S8). HMR isolates were subsequently tested for growth on PYT80 with ampicillin, kanamycin, or erythromycin to determine prevalence of cross-resistance (Groves et al., 1975; Wang et al., 2021); strains forming visible colonies in the presence of antibiotics within 7 days were considered tolerant.

## RESULTS

### Impact of HM pollution on soil bacterial community structure

Soils from sampling sites in 35207 had significantly higher levels of Pb, Mn, and Zn than those in 35214 (Mann-Whitney U-tests, *p-value < 0*.*05*). As and Cd were below detection limits for all samples with the exception of 35207 site F having 37 ppm As. Average silhouette width k-mer clustering based on metal concentrations supported clustering the samples into two groups that corresponded exactly to the two neighborhood zip codes, supporting our hypothesis that 35207 soils were significantly more exposed to HM than 35214 (Fig. 1B). Despite the clear evidence of elevated metals in 35207, only Mn (in all 12 samples) and As (in one 35207 sample) concentrations exceeded EPA Residential Soil Regional Screening Level (RSL) guidelines for human health concerns (As 0.68, Cd 7.1, Pb 400, Mn 180, and Zn 2300 ppm) (Li et al., 2019, Li, 2018).

Both NMDS and CCA ordination techniques revealed significant correlations of Mn concentration and pH on soil community structure (Figs. 2A, S1A; NMDS ordination also showed a significant effect of Zn). NMDS ordination did not show a significant clustering of samples by zip code (Fig. S1A, AMOVA p > 0.05), but when ordination was constrained using soil metadata (HM concentrations, pH, and organic carbon content) using CCA, the zip codes were clearly differentiated (Fig. 2A). While pH was a significant structuring force, only metal concentrations were important for separating the zip codes, with all 4 metal biplot vectors indicating movement toward the upper right corner of the plot (Fig. 2A).

**Figure 2.**
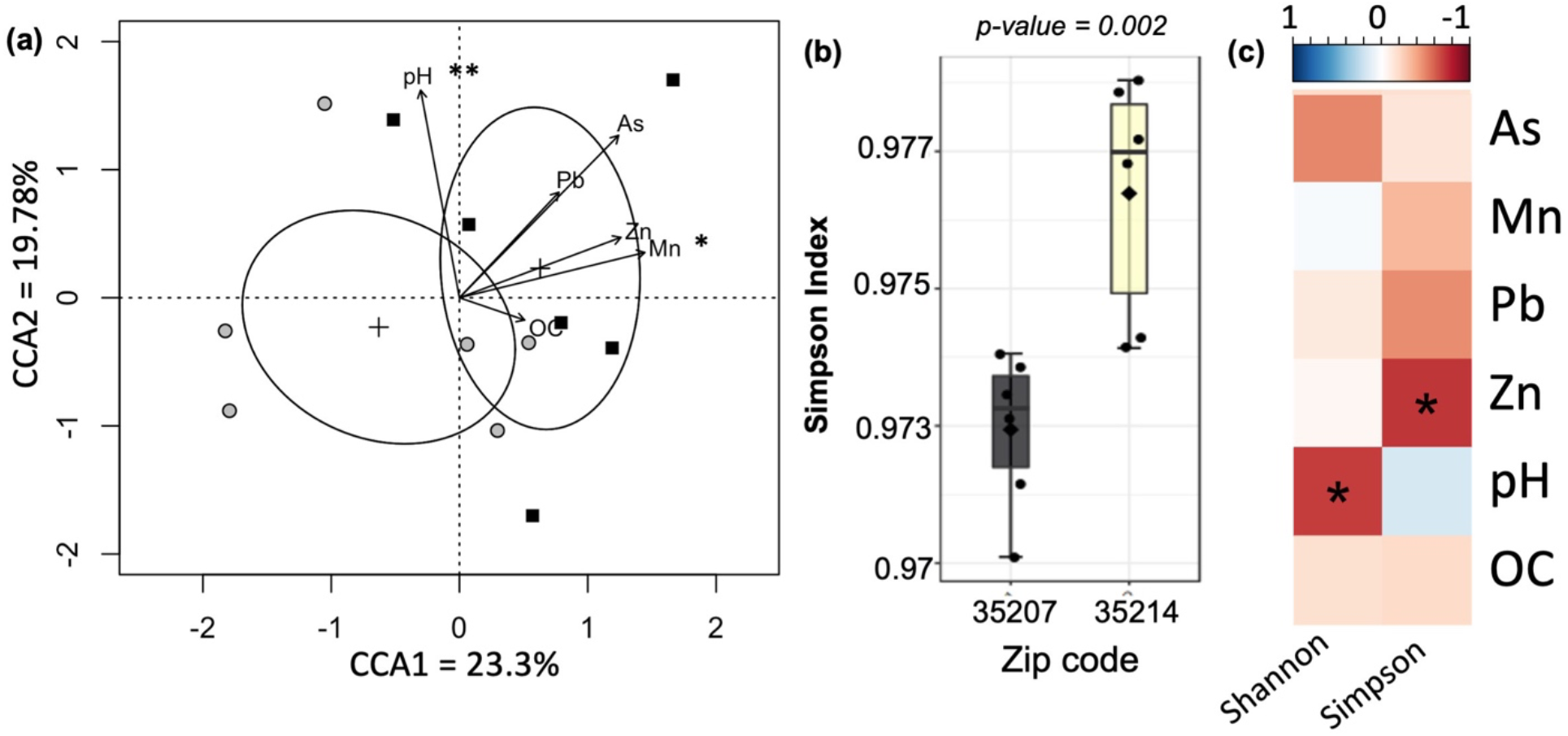
**(a)** Multivariate canonical correspondence analysis (CCA), the ellipses represent 95% confidence intervals of the centroid of the groups and the vectors indicate the impact of metal concentrations and other environmental variables on the position of points in the plot.; **(b)** Alpha diversity of bacterial community at genus level (Simpson Index). **(c)** Spearman correlation between metal concentrations and the indicated alpha diversity metrics.

<u>There was also some evidence of a negative impact of metals on soil bacterial diversity. The</u> Simpson alpha diversity index, calculated based on OTUs clustered phylogenetically at the genus level, was significantly different between zip codes (Fig. 2B). However, the Shannon index calculated the same way did not show a significant difference, nor did either metric calculated at the OTU level without phylogenetic clustering. Interestingly, both metrics were correlated with metadata; the Shannon index indicated that diversity decreased with increasing pH, and the Simpson index was negatively correlated with Zn concentration (Fig. 2C).

### Impact of metals on bacterial taxonomic groups

Across all samples, the most abundant bacterial phyla were *Actinobacteriota, Acidobacteriota, Proteobacteria*, and *Chloroflexi*, representing 73%-83% of total bacteria (Fig. 3A). Nine of the 48 OTUs that reached a relative abundance of at least 1% in at least one sample were significantly different between sites (Fig. 3B). Of these 9, 6 were more abundant in 35207, including the most abundant of the 9, a representative of the gram-positive Solirubrobacterales 67-14 clade. Of higher-level taxa comprising at least 1% of one sample, 2 phyla, 4 classes, 5 orders, 5 families, 6 genera, and 1 species were significantly different between sites (Fig. 3C). Notably, the highly abundant phylum Proteobacteria was significantly less abundant in 35207, whereas Methylomirabilota, a poorly studied group containing the as-yet uncultivated Rokubacteriales, was more abundant in 35207. While the phyla Acidobacteriota and Actinobacteriota were not overall different between zip codes, specific subgroups (e.g. the Blastocatellia and Solirubrobacterales, both more abundant in 35207) were.

**Figure 3.**
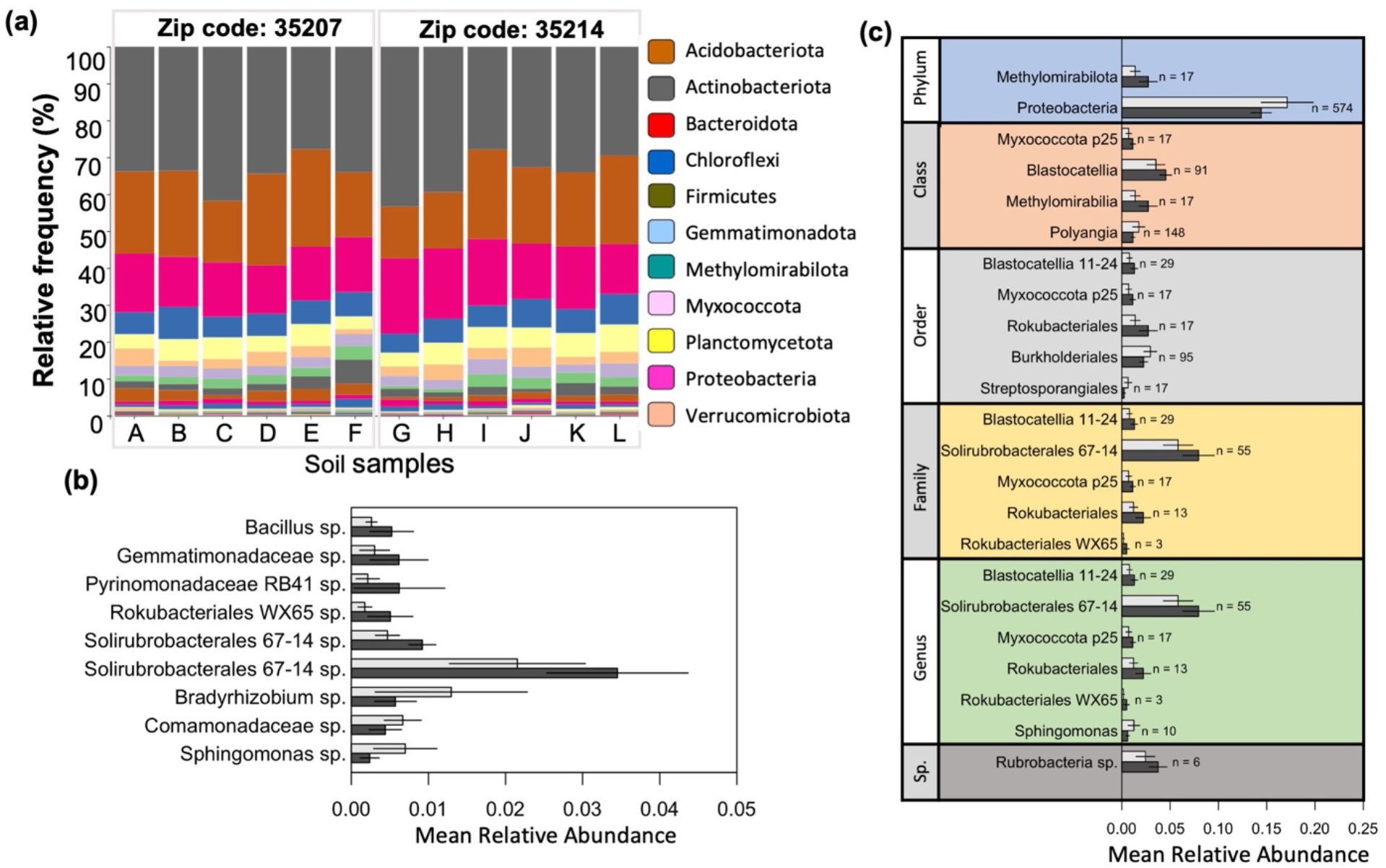
**(a)** Relative abundance of phyla of soil bacterial community from 16s rRNA sequencing. The taxonomy bar plot only highlights phylum names which were at least 1% or more relatively abundant; **(b-c)** Relative abundances of OTUs **(b)** or higher-level taxa **(c)** that were significantly different between 35207 (dark gray bars) and 35214 (light gray bars). Only taxa that were at lease 1% of at least one community are shown.

A number of taxa were significantly correlated with environmental parameters measured at the sampling sites, including metals (Figures S2-S4). Higher organic carbon content was correlated with higher ratios of archaea to eubacteria, driven by the abundance of OTUs similar to the ammonia-oxidizing archaeal genus *Nitrosphaera*. Lower pH was correlated with greater representation of bacteria from the phylum Planctomycetota. Two classes within the phylum Chloroflexi were also correlated with pH, with the Chloroflexia favoring low pH and the KD4-96 clade favoring higher pH. Unsurprisingly, the Acidobacteriota family Vicinamibacteraceae was correlated with lower pH.

At the phylum level, only the Verrucomicrobiota were correlated with metal concentrations, being rarer in higher Mn samples. Class Polyangia (Myxococcota) as well as Orders Thermoanaerobaculales (Acidobacteriota) and Streptosporangiales (Actinobacteriota) were negatively correlated with Zn, and Tepidisphaerales (Planctomycetota) was negatively correlated with both Zn and Pb. Other taxa were positively correlated with metals: the Myxococcota bacteriap25 class was more abundant in higher Zn samples, the Solirubrobacterales 67-14 clade was positively associated with both Pb and Zn, and the Rokubacterales WX65 genus was positively correlated with both Mn and Zn. No significant correlations were observed between any taxon and As.

Interestingly, of the 48 OTUs that comprised at least 1% of at least 1 sample, no significantly negative correlations with metal concentrations were observed, whereas 11, or 23%, were positively correlated with at least one metal (Fig. S4). Of these, 5 were positively associated with 2 metals, and 3 with all three of Mn, Pb, and Zn. 6 of these OTUs were identified as significantly different between zip codes by both Metastats and LEFSe analyses; of these, 5 were positively correlated with at least 2 metals. 16 of the 48 (33%) were also significantly correlated with at least one of the first two NMDS axes (Fig. S1B), with three biplot vectors (Pyrinomonadaceae RB41, Rubrobacteria sp., and Gemmatimonadaceae sp., all also positively correlated with metals) pointing into the same quadrant as the metal biplot vectors.

### Influence of metals on predicted soil metagenomes

We used PICRUSt2 to infer the metagenomes of our 12 soil samples based on 16S profiles, predicting 7637 unique KO IDs which allowed us to predict the functional potential of the microbial communities. The functional profiles of the two zip codes were structured significantly differently (NMDS on Bray-Curtis distance, non-overlapping 95% confidence intervals of centroids, Fig. S5B), with much greater dispersion in the ordination coordinates of the 35214 samples than those from 35207 (areas of the 95% confidence interval ellipsoids 0.011 and 0.003 respectively). The same general conclusions held when we constrained the ordination to just the AMR and HMR genes in our predictions (Fig S5B, ellipsoid areas of 35214 and 35207 0.013 and 0.006 respectively). We found that genes from two-component sensory systems, carbon fixation and catabolism pathways, and vancomycin resistance were most likely to differ between the zip codes (Fig. 4a).

**Figure 4.**
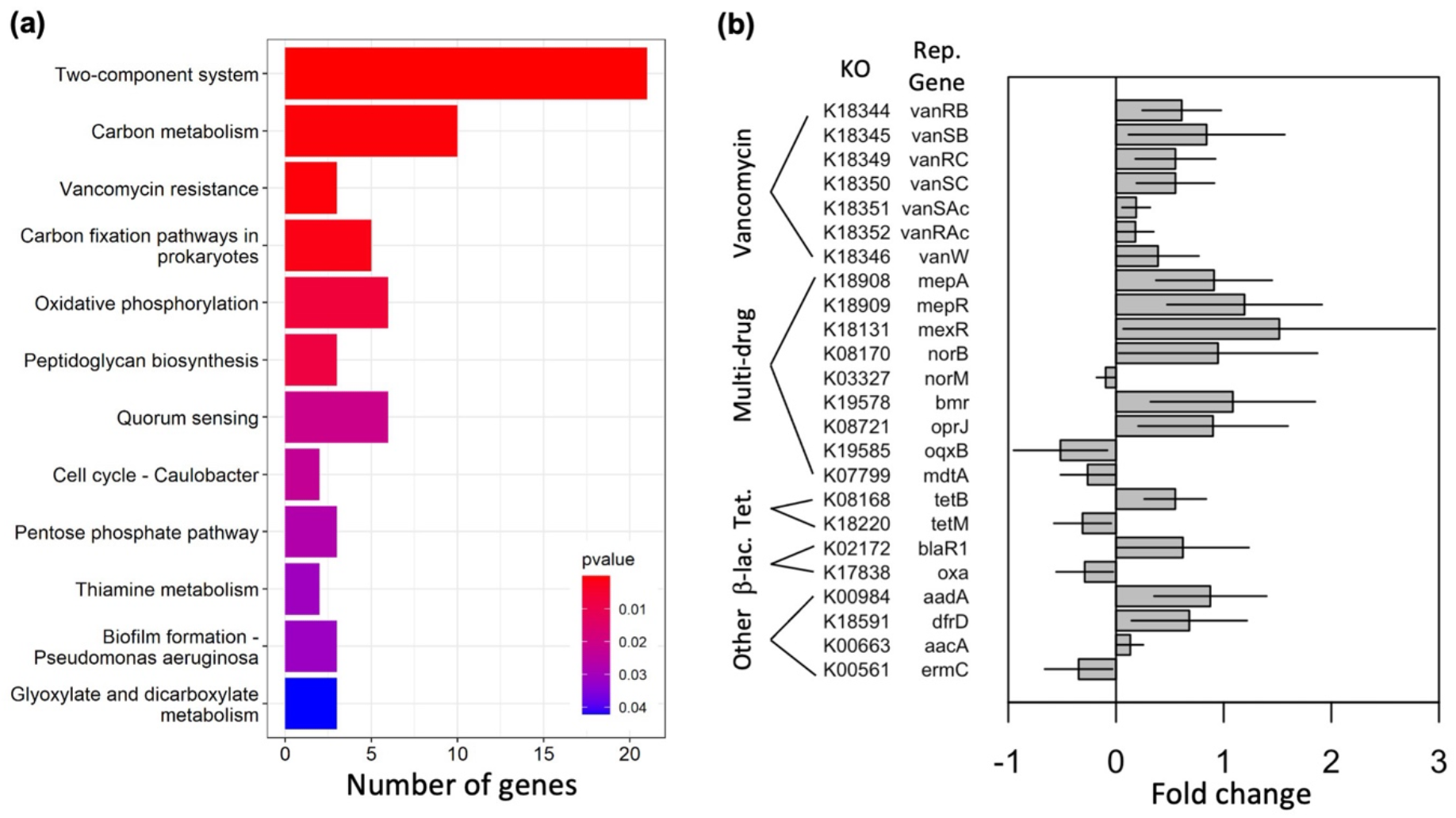
**(a)** Over-representation analysis of differences between gene presence/absence in 35207 vs. 35214 based on PICRUSt metagenome inference. Bars indicate the number of genes found to significantly differ between the zip codes that fall into the indicated pathway; p-values indicate the probability of finding that many differentially represented genes by chance. **(b)** Antimicrobial genes that differ significantly between zip codes based on PICRUSt metagenome inference. Positive values indicate enrichment in 35207; negative values indicate enrichment in 35214. Error bars represent the 95% confidence interval of the estimate.

We assessed the difference between predicted gene abundances in the two zip codes using Welch’s t-test in STAMP. Out of 7637 genes, 377 were differentially abundant between the two zip codes (adjusted *p-value* < 5%, Table S6). 24 of these genes, or 6.4%, were associated with AMR pathways, and 9 were associated with HMR (Fig. 4B, Table S9). Overall, 46.2% of the significantly differently abundant genes were more abundant in 35207 samples, compared to 75% of differently abundant AMR genes; AMR genes were thus significantly more likely to be different between the sites based on a Fisher’s exact test (p = 0.005). All 7 identified genes related to vancomycin resistance were more abundant in 35207, as well as 6 of 9 genes identified as being involved in multidrug resistance. Interestingly, only 5 of 9 genes related to HMR were significantly more abundant in 35207, and HMR genes were not more likely to be significantly different between zip codes than other types of genes (Fisher’s exact test, p = 0.41).

We also observed direct correlations between metal concentrations and the abundances of predicted AMR and HMR genes. Altogether, Pb, Mn, and/or Zn were significant predictors of 21 AMR/HMR gene abundances (linear model, p < 0.05 for the slope of metal concentration vs. abundance being not equal to zero, Table S10). 100% of the genes significantly impacted by Pb were more abundant at higher Pb concentrations, and only 1 of 12 genes impacted by Mn was less abundant at higher Mn concentrations (Table 1). Zn, on the other hand, was negatively related to AMR/HMR gene abundance in 5 of 6 significant interactions, and in all 5 cases genes that were negatively related to Zn concentration were positively related to Pb concentration.

**Table 1.** AMR and HMR genes predicted by metal concentrations. +, positive slope of gene abundance vs. metal concentration; −, negative slope

| Class | KO | gene | annotation | Pb | Mn | Zn |
| --- | --- | --- | --- | --- | --- | --- |
| AMR | K00561 | ermC,<br>ermA | 23S rRNA (adenine-N6)-dimethyltransferase [EC:2.1.1.184] | + |  | - |
| AMR | K08217 | mef | MFS transporter, DHA3 family, macrolide efflux protein | + |  | - |
| AMR | K18220 | tetM,<br>tetO | ribosomal protection tetracycline resistance protein | + |  | - |
| AMR | K08167 | smvA,<br>qacA, lfrA | MFS transporter, DHA2 family, multidrug resistance protein | + |  |  |
| AMR | K19062 | arr | rifampin ADP-ribosylating transferase | + |  |  |
| AMR | K18909 | mepR | MarR family transcriptional regulator, repressor for mepA |  | + | + |
| AMR | K18131 | mexR | MarR family transcriptional regulator, repressor of the mexAB-oprM multidrug resistance operon |  | + |  |
| AMR | K18589 | dfrA1,<br>dhfr | dihydrofolate reductase (trimethoprim resistance protein) [EC:1.5.1.3] |  | + |  |
| AMR | K18780 | blaNDM | metallo-beta-lactamase class B NDM [EC:3.5.2.6] |  | + |  |
| AMR | K18792 | blaOXA-10 | beta-lactamase class D OXA-10 [EC:3.5.2.6] |  | + |  |
| AMR | K18824 | sul2 | dihydropteroate synthase type 2 [EC:2.5.1.15] |  | + |  |
| AMR | K19096 | blaCMY-2 | beta-lactamase class C CMY-2 [EC:3.5.2.6] |  | + |  |
| AMR | K19274 | aph3-VI | aminoglycoside 3'-phosphotransferase VI [EC:2.7.1.95] |  | + |  |
| AMR | K19278 | aac6-Ib | aminoglycoside 6'-N-acetyltransferase Ib [EC:2.3.1.82] |  | + |  |
| AMR | K19276 | aac3-IV | aminoglycoside 3-N-acetyltransferase IV [EC:2.3.1.81] |  | - |  |
| Metal | K07156 | copC,<br>pcoC | copper resistance protein C | + |  | - |
| Metal | K07241 | nixA | high-affinity nickel-transport protein | + |  | - |
| Metal | K07665 | cusR,<br>copR, silR | two-component system, OmpR family, copper resistance phosphate regulon response regulator CusR | + |  |  |
| Metal | K07230 | p19, ftrA | periplasmic iron binding protein | + |  |  |
| Metal | K07311 | ynfG | Tat-targeted selenate reductase subunit YnfG |  | + |  |
| Metal | K07490 | feoC | ferrous iron transport protein C |  | + |  |

### Cross-resistance to antimicrobials in HMR bacterial isolates

We tested the ability of bacteria from each sample to grow on media containing metals or antibiotics. There were no significant differences in AMR or HMR between samples in 35214 and 35207 when intact soil communities were diluted onto agar plates (Wilcoxon tests, p > 0.05, Fig. 5A), nor were samples from sites with higher metal concentrations significantly more AMR or HMR (linear models, p > 0.05). With the exception of Pb (in both zip codes) and ampicillin (in 35207), all additions significantly reduced bacterial growth relative to the unamended control plates (Wilcoxon signed rank tests, p < 0.05).

**Figure 5.**
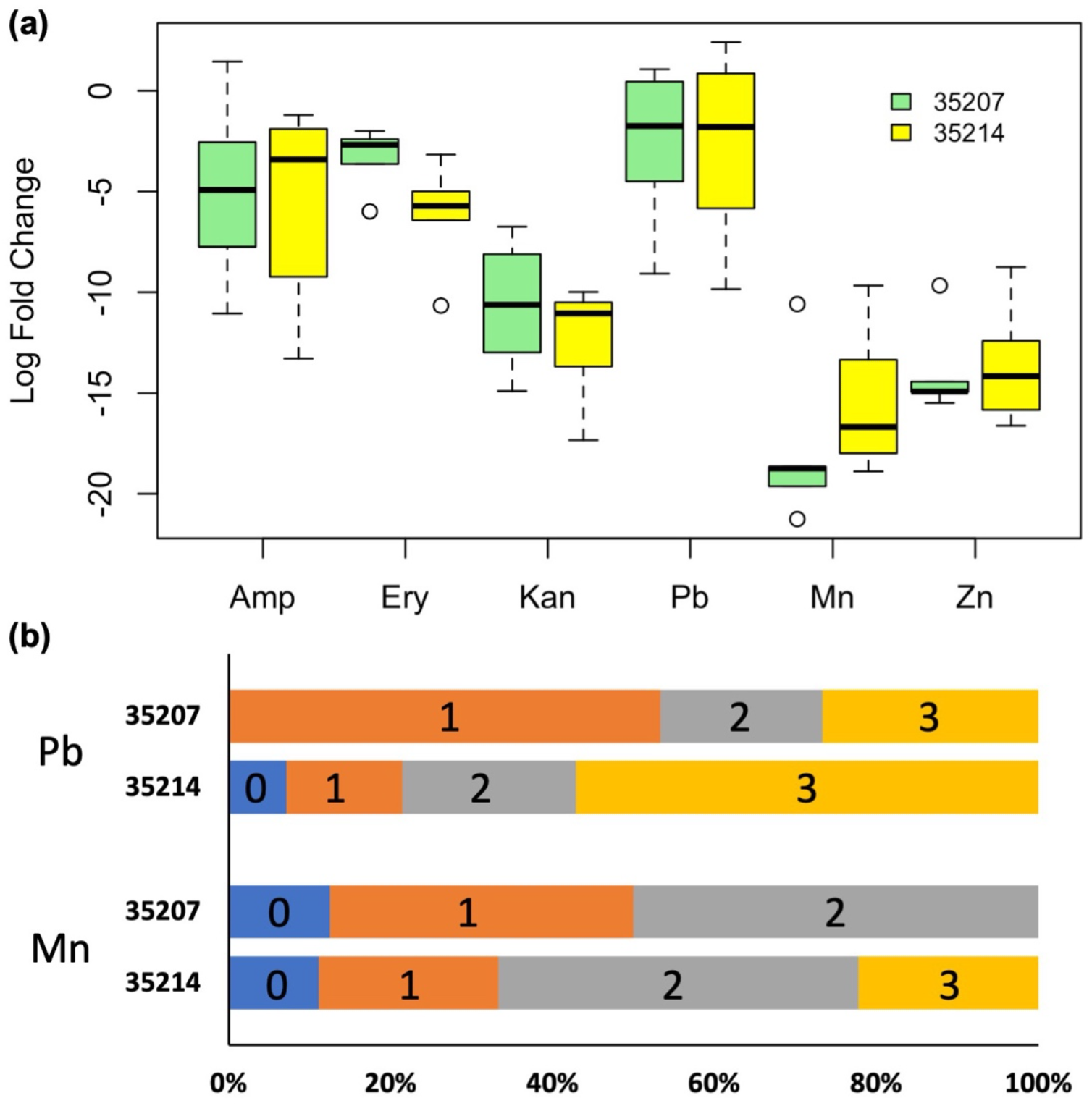
**(a)** Recovery of colonies on antibiotic and heavy metal treated PYT80 plates from soil samples taken from zip codes 35207 and 35214, shown in green yellow boxes respectively. Values indicate the base 2 logarithm of the ratio of growth on metal-or antibiotic-treated plates to growth on unamended plates. Bars represent the interquartile range, with the central bar representing the median value and the whiskers extending to the most extreme points equal to or less than 1.5 times the interquartile range from the median, with circles representing outliers beyond this range. **(b)** Bars indicate the proportion of bacterial isolates taken from plates containing the indicated metal that were also resistant to the indicated number of the 3 tested antibiotics (ampicillin, erythromycin, and kanamaycin).

To assess the resistance phenotypes of individual strains from each zip code, we isolated 46 strains across zip codes that were able to grow on PYT80-HM plates spiked with Mn (17 isolates) or Pb (29 isolates); no isolates were obtained from Zn-spiked plates. Based on 16S sequences, 24 taxonomically distinct isolates were identified based on their closest match in the BLAST nr database (Fig. S6). These isolates were heavily concentrated in the phyla Actinobacteriota and Proteobacteria and were not broadly representative of the taxonomic diversity revealed by our 16S tag sequencing efforts (Fig. 3). 12 of the bacterial isolates were unique to 35207, 8 were unique to 35214, and 4 were common across both zip codes. We then tested each of the 46 strains’ tolerance to PYT80 plates supplemented with ampicillin, kanamycin, or erythromycin. In contrast to the broad inhibition of the bulk communities by antibiotics, 94% of HMR isolates were resistant to at least one antibiotic, and 30% were resistant to all 3 (Fig. 5B). There was no significant difference between zip codes or the metal spike used during strain isolation in the number of antibiotics a strain was resistant to (linear mixed effects model with phylogenetic identification as the random effect, p > 0.05 for each fixed effect predictor) or in the likelihood that a strain was specifically resistant to ampicillin or kanamycin (binomial mixed effects regression, p > 0.05 for each predictor). Strains isolated from 35214 were, however, significantly more likely to be resistant to erythromycin (binomial mixed effects regression, p = 0.02). For taxa that were isolated multiple times, there was substantial variation between strains in the antibiotic resistance profile (Table S8). For instance, out of 9 *Rhodococcus degradans* isolates, 6 were resistant to all antibiotics, 2 were susceptible to erythromycin, and one was susceptible to all three antibiotics tested.

## DISCUSSION

The 35^th^ Avenue Superfund site in North Birmingham, Alabama, houses coke and coal industries that have left a legacy of HM pollution behind, increasing the risk of lung and other diseases for local residents (Allen et al., 2019; GASP, 2014). Ongoing efforts to understand and mitigate the human impacts of this environmental injustice have generally looked to the direct effects of HMs on human biology, but to our knowledge there is little work exploring the possible relationship between HM contamination and AR prevalence at a US Superfund site.

First, we asked simply whether soil bacterial communities from the 35^th^ Avenue Superfund site differed from those in a less polluted nearby neighborhood. HMs As, Pb, Mn and Zn were all more abundant in 35207 compared to 35214, and kmer clustering confirmed that the zip codes could be distinguished by their metal profiles (Fig. 1B). While only Mn and As ever surpassed EPA guidelines determining unacceptable levels of residential soil pollution, HMs nevertheless had a significant effect on microbial community composition (Fig 2A, Fig S1A). Metal contaminated sites in 35207 were significantly less diverse than 35214 (Fig. 2B), with Zn having an especially negative impact on overall diversity (Fig. 2C).

The abundance of a number of bacterial taxa were significantly affected by chronic metal polluted soils (Fig. 3). Some taxa were significantly different between the zip codes, including highly abundant groups such as the Proteobacteria (lower in 35207) and the Solirubrobacterales (higher in 35207). Absolute metal concentrations were also significant predictors of taxon abundance, and approximately 1 in 4 of the most abundant OTUs in our dataset were positively correlated with at least one of the metals Mn, Pb, and/or Zn (Fig. S4). Overall, the magnitude of impact of metals on community structure was roughly equivalent to that of pH, which has been shown previously to be the dominant structuring factor for soil microbiomes across many diverse ecosystems (Fierer and Jackson 2006, Lauber, Hamady et al. 2009).

Our hypothesis that AMR was more common in metal-contaminated soils in our study site was supported by our inferred metagenomes, where numerous genes related to antimicrobial resistance were significantly more abundant in 35207 (Fig. 4) and/or positively correlated with metal concentration. We also found that bacterial isolates from our soil samples that were selected using HM spiked agar were also highly likely to be resistant to multiple antibiotics (Fig. 5B). However, we were unable to detect a significant difference in community-scale phenotypic AMR or HMR in soil bacterial communities (Fig. 5A). It is possible that this discrepancy reflects biases in our metagenome inference software, as the NSTI values for our samples indicated relatively low representation of many of our taxa in published databases (Langille, Zaneveld et al. 2013), although our pipeline removed highly divergent OTUs from the metagenome inference to minimize this problem. A more likely cause is that culture-based assays of soil communities are potentially misleading due to the strong cultivation bias in these systems (Harwani, 2013). The great majority of our isolates fell into a few clades that were not closely related to the most abundant taxa from our tag sequencing analysis (Fig. S6), and it is noteworthy that only three of the 48 OTUs comprising more than 1% of any sample (*Bacillus, Rhizobium*, and *Streptomyces*) had a close relative amongst the isolates.

Our inferred metagenomes discovered AMR genes related to a wide variety of antibiotic targets such as protein synthesis (tetracycline resistance), peptidoglycan synthesis (beta-lactam and vancomycin resistance), and folate synthesis (trimethoprim resistance), nearly all of which were significantly higher in samples from 35207 (Fig. 4B). Both vancomycin and trimethoprim resistance have been previously shown to co-occur with resistance against various HMs (Zhong et al., 2021; Dickenson et al., 2019). We also discovered a number of multi-drug resistance genes such as efflux pumps enriched in 35207 samples. Importantly, these genes include many of the most clinically concerning AMR pathways and reinforce the importance of considering the impact of environmental pollution as an additional vector for AMR evolution, along with clinical and agricultural antibiotic use.

In conclusion, our study provides tentative support for the hypothesis that chronic heavy metal pollution in soils may select for antimicrobial resistance that may negatively impact humans living near those soils, reflecting an additional health concern beyond those caused by the HMs themselves. However, our results fall short of demonstrating a strong connection between HMR and AR at the 35^th^ Avenue Superfund site due to methodological limitations. For example, PICRUSt2 accuracy is higher for human microbiome samples than for soils (Langille et al., 2013) so our prediction results should be cautiously interpreted, but also even under ideal circumstances it is likely difficult to infer the presence of highly mobile genes like those involved in AMR merely from the taxonomic information provided by 16S tag sequencing. Future work should target specific HM or AMR genes of interest using quantitative PCR (Lin et al., 2012) or plasmidomics/mobilomics to improve detection of resistance genes on mobile genetic elements that may be exchanged between soil microbial species and possibly between those organisms and counterparts in the human microbiome (Schlüter et al., 2008; Li et al., 2015). Culture-based assays could also be improved, for instance by performing 16S tag sequencing of enrichment cultures following exposure of intact communities to selective concentrations of HMs or antibiotics. Future work should also incorporate more detailed datasets and ideally should also involve the human residents of the zip codes. Ultimately, being able to connect the environmental microbiome to the human microbiomes of 35207 residents could help determine the degree to which HMs impact AR diagnoses like recalcitrant infections, and could help improve health care for this vulnerable population.

## ACKNOWLEDGEMENTS

We thank Dr. Rob Akscyn as well as instructors and students across our multiple course-based undergraduate experiences for preliminary methods troubleshooting. We are grateful to Dr. Veena Antony for leading us to this work. We thank Dr. Dustin Kemp for access to equipment, and Dr. Casey Morrow and the UAB Genomics Core for sequencing services. Lastly, Dr. Shauntice Allen and grassroots organizations such as GASP and PANIC continue to show us what it means to advocate for the North Birmingham community.

## AUTHORS’ CONTRIBUTION

<u>Anuradha Goswami</u>* - Conceptualization, Analysis, Writing and editing *Indicates co-first authorship

<u>Sarah J. Adkins-Jablonsky</u>* Methodology (Soil sampling, 16S DNA extraction, culture assays and preparation), Funding acquisition, writing and editing *Indicates co-first authorship

<u>Marcelo Malisano Barreto Filho – gene overrepresentation analysis</u>

<u>Michelle D. Curtis, Alex Dawson, Sabrina Heiser, Aisha O’Connor, Melissa Walker-</u>Analysis (16S sequence processing using MOTHUR), Writing-editing and review

<u>Qutia Roberts</u> - Methodology (Soil sampling, 16S DNA extraction)

<u>J. Jeffrey Morris</u> - Investigation and conceptualization, Supervision, Analysis, Funding acquisition, Writing and editing

