## Supplementary material for "Heavy metal pollution impacts soil bacterial community structure and antimicrobial resistance at the Birmingham 35^th^ Avenue Superfund Site": Fig. S

**Supplemental Figures for Goswami et al. 2022**

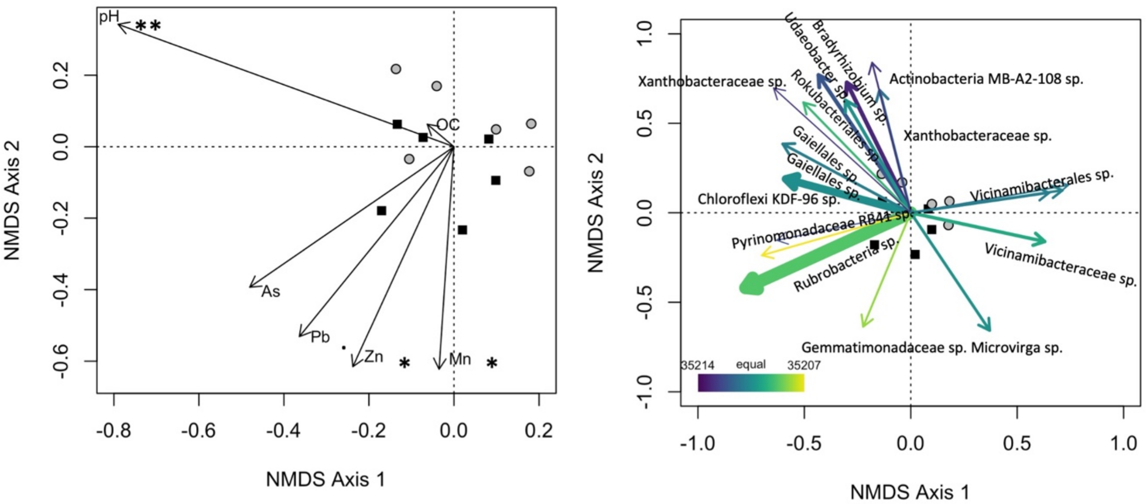


**Figure S1. NMDS ordination plots of soil microbial communities. (a)** Biplot vectors showing the influence of soil metadata, including metal concentrations, pH and organic carbon (OC) content on position of communities in the plot. Asterisks indicate the significance value of the correlation of the most significant axis with the indicated parameter; *, $p < 0.05$; **, $p < 0.01$. **(b)** The same plot, but showing the biplot vectors of all OTUs that represented at least 1% of any sample. The width of the arrow is proportional to the average relative abundance of the OTU across all samples, and the color indicates the level of bias towards one zip code or the other. Black squares = 35207 samples; gray circles = 35214 samples.

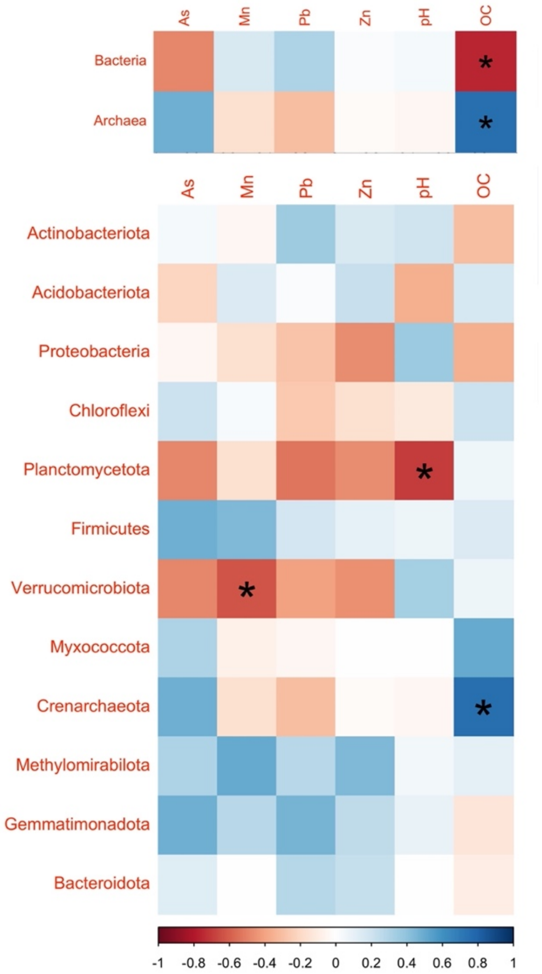


**Figure S2. Correlation of Domain and Phylum taxa with soil metadata.** Color scale reflects the Spearman correlation coefficient of the taxon's abundance vs. the indicated metadata parameter from each soil sample. *, $p < 0.05$.

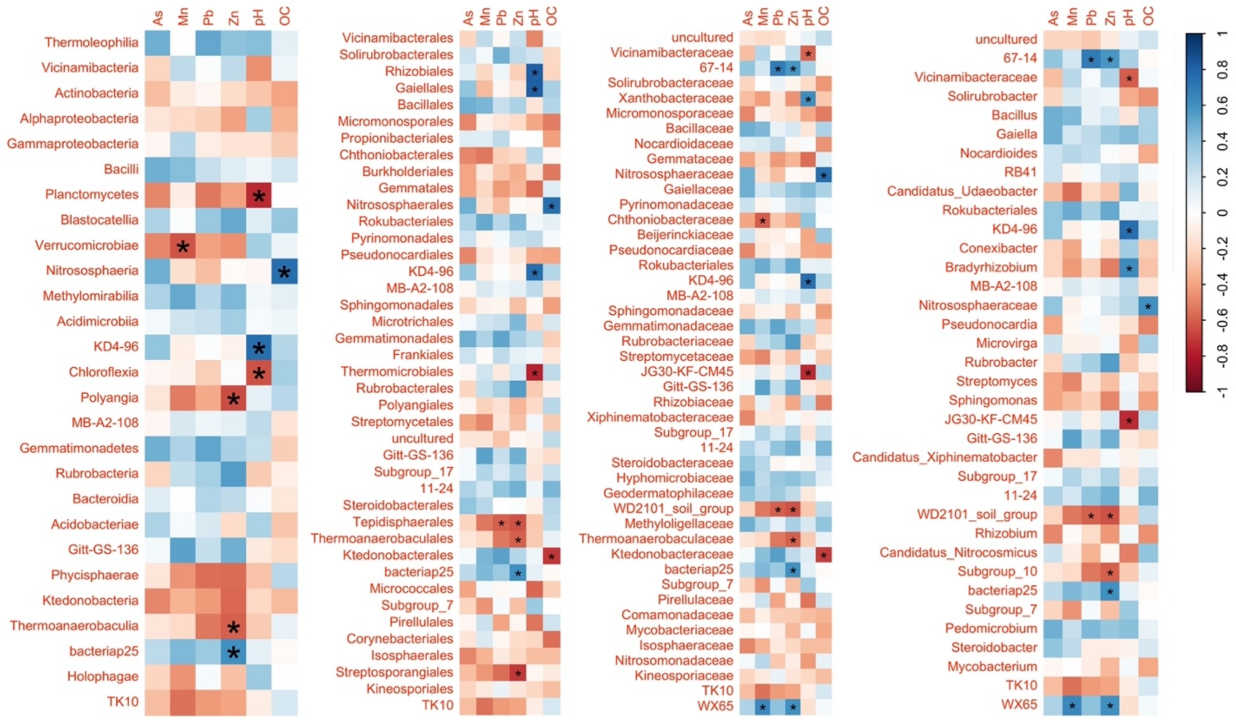


**Figure S3. Correlation of Class, Order, Family, and Genus taxa with soil metadata.** Color scale reflects the Spearman correlation coefficient of the taxon's abundance vs. the indicated metadata parameter from each soil sample. *, $p < 0.05$.

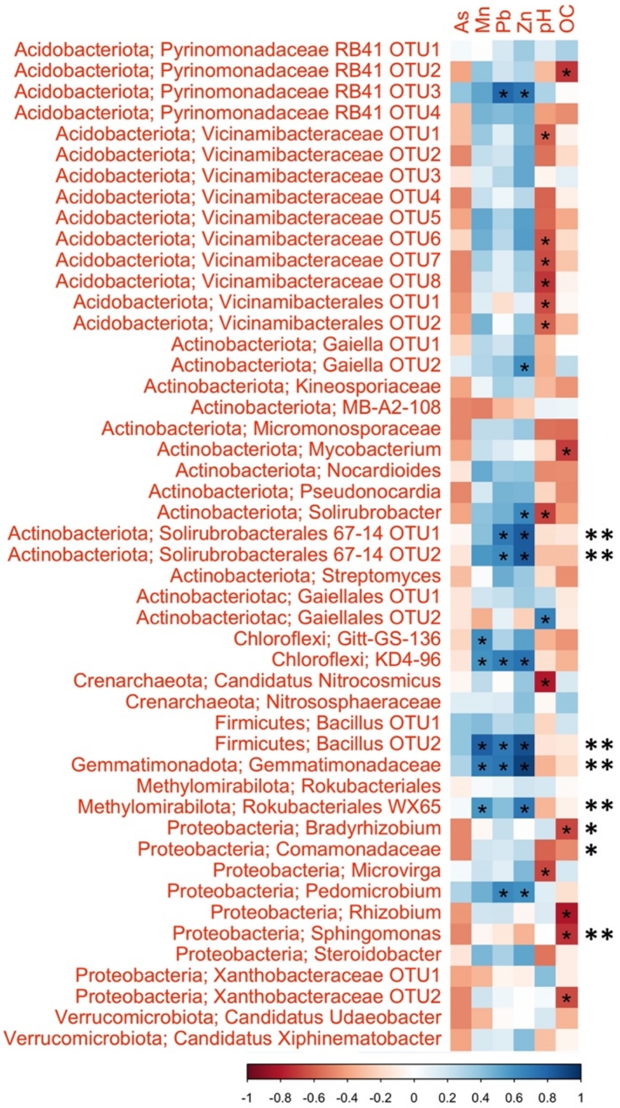


**Figure S4. Correlation of OTUs with soil metadata.** All OTUs representing at least 1% of any soil sample are shown, along with their taxonomic identifications at the phylum and genus levels. Color scale reflects the Spearman correlation coefficient of the taxon's abundance vs. the indicated metadata parameter from each soil sample. Asterisks within colored squares indicate the significance level of the correlation coefficient, with * indicating $p < 0.05$. Asterisks to the right of the heat map indicate taxa that returned significant adjusted p-values in Metastats (one asterisk) and/or LEFSe (two asterisks) analyses.

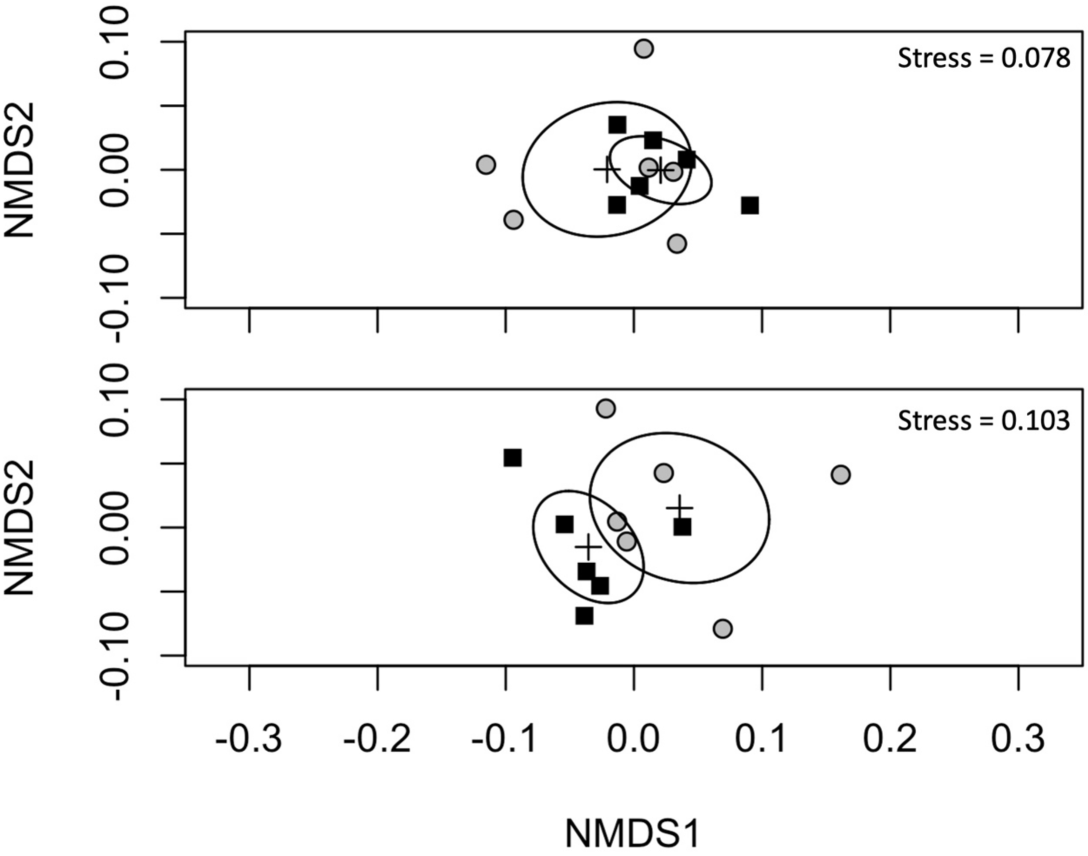


**Figure S5. NMDS ordination plots of inferred gene abundance from PICRUSt analysis.** The ordination in the top plot used all predicted gene abundances, whereas the bottom plot was restricted to AMR and HMR genes. Crosses and ellipses indicate the centroids and 95% confidence intervals of the zip codes. Black squares = 35207 samples; gray circles = 35214 samples.

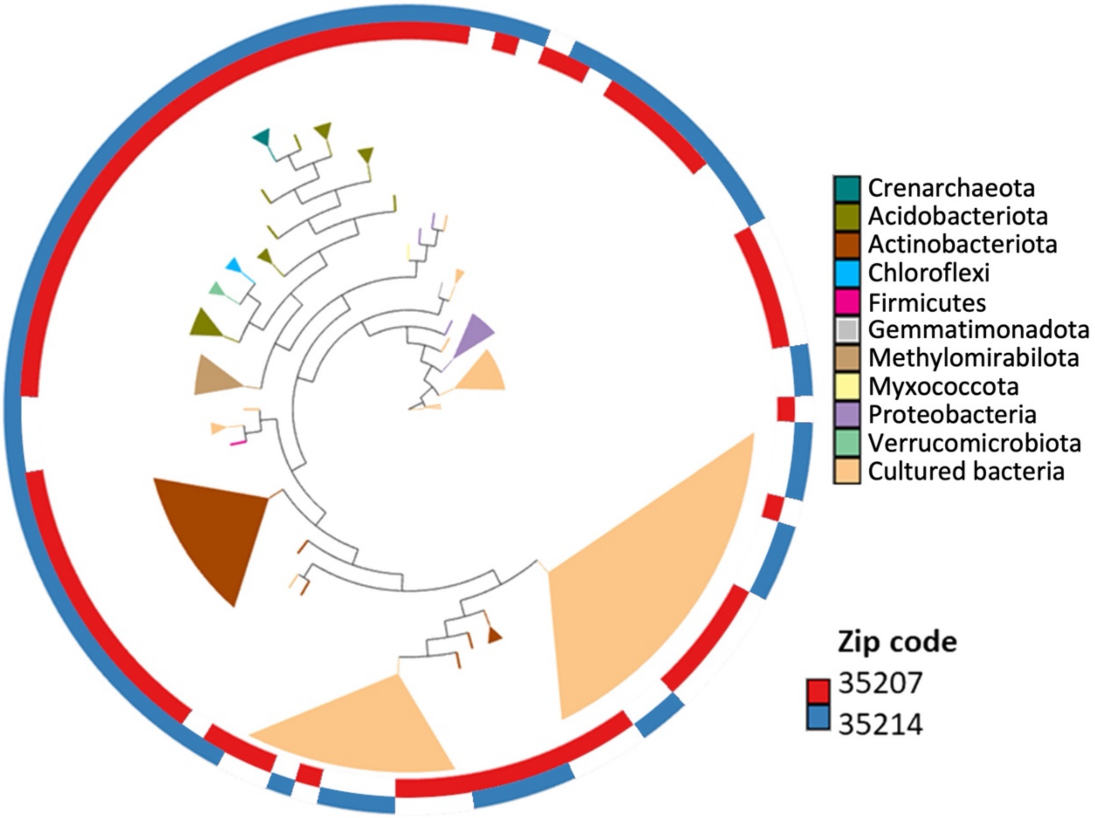


**Figure S6. Phylogenetic tree of most abundant OTUs and cultured isolates used in HMR and AMR assays.** The phylogenetic tree was assembled using the 16S rRNA sequences of all OTUs representing at least 1% of any sample as well as the 16S samples of all isolates used in our culture-based assays. Colored wedges represent multiple sequences with the same phylum-level taxonomic assignment within the same clade. Inner and outer rings represent tax found in 35207 and 35214.
